# The repetitive DNA sequence landscape and DNA methylation in chromosomes of an apomictic tropical forage grass, *Cenchrus ciliaris*

**DOI:** 10.1101/2022.05.25.493456

**Authors:** Priyanka Rathore, Trude Schwarzacher, J.S. (Pat) Heslop-Harrison, Vishnu Bhat, Paulina Tomaszewska

## Abstract

*Cenchrus ciliaris* is an apomictic, allotetraploid pasture grass widely distributed in tropical and subtropical regions of Africa and Asia. In this work, we aim to investigate the genomic organization and characterize the nature of repetitive DNA sequences in this species. Because of the apomictic propagation, various aneuploid genotypes are found and we analysed here a 2*n*=4*×*+3=39 accession. The physical mapping of Ty1-*copia* and Ty3-*gypsy* retroelements through fluorescence *in situ* hybridization with global assessment of 5-methylcytosine DNA methylation through immunostaining revealed the genome-wide distribution pattern of retroelements and their association with DNA methylation. About a third of Ty1-*copia* sites overlapped or spanned centromeric DAPI positive heterochromatin, while the centromeric regions and arms of some chromosomes were labeled with Ty3-*gypsy*. Most of the retroelement sites overlapped with 5-methycytosine signals, except some Ty3-*gypsy* on the arms of chromosomes which did not overlap with anti-5-mC signals. Universal retrotransposon probes did not distinguish genomes of *C. ciliaris* showing signals in pericentromeric regions of all 39 chromosomes, unlike highly abundant repetitive DNA motifs found in survey genome sequences of *C. ciliaris* using graph-based clustering. Probes developed from RepeatExplorer clusters gave strong signals mostly in pericentromeric regions of about half of the chromosomes, and we suggested that they differentiate the two ancestral genomes in the allotetraploid *C. ciliaris* likely having different repeat sequence variants amplified before the genome came together in the tetraploid.

## Introduction

*Cenchrus ciliaris* L. (buffelgrass, buffel-grass, or African foxtail grass; syn. *Pennisetum ciliare*) is an apomictic, allotetraploid perennial C_4_ grass native to tropical and subtropical arid regions of Africa and Western Asia. Predominantly distributed in warmer tropical regions, *C. ciliaris* grows in a range of harsh conditions (Marshall et al., 2012; Kharrat-Souissi et al., 2013, 2014). *Cenchrus* species are the predominant pasture grasses in India due to their high fodder value (Meena and Nagar, 2019). They are of great ecological importance due to persistence in native arid ecosystems, associated with tolerance to heavy grazing and drought, with deep root systems, and they are also used for erosion control (Miller et al., 2010). *C. ciliaris* is recognized as an invasive species in Australia (where it is a ‘declared plant’), South America and the USA (particularly in Arizona and the Sonoran Desert) and became a threat to natural ecosystems (Jackson, 2005; Miller et al., 2010) both by displacement of native plants and by intensifying wildfires. It is important to study the genetic characteristics and genomic organization of *C. ciliaris* to understand the association between biological, agricultural, and ecological characteristics.

*Cenchrus ciliaris*, with a base chromosome number of *×*=9, includes tetra-, penta-, hexa- and septa-ploid cytotypes (Visser et al., 1998; Kharrat-Souissi et al., 2013; Dhaliwal et al., 2018), and aneuploid accessions with 2*n*=32, 40, 43, 44 and 48 have also been found (Burson et al., 2012; Carloni-Jarrys et al., 2018). Most accessions are apomictic, although sexual plants are also found occasionally in nature (Fisher et al., 1954; Yadav et al., 2012; Marshall et al., 2012), with a DNA-based SCAR marker reported by Dwivedi et al. (2007). Apomictic seed production preserves superior genotypes, including heterozygotes and polyploids (Spillane et al., 2004; Bhat et al., 2005; Abdi et al., 2016) and can found new polyploid lineages. The character is well represented in polyploid grasses including members of Paniceae grass genera such as *Panicum, Pennisetum, Cenchrus* and *Paspalum* (Worthington et al., 2016; Higgins et al., 2021; Tomaszewska et al., 2021b). In *Urochloa* (*Brachiaria*), there is a robust molecular marker discriminating sexual and apomictic plants. The genes involved in *C. ciliaris* have been examined and compared with other species (Roche et al., 1999; Akiyama et al., 2005; Yadav et al., 2012). Like the widely grown *Urochloa* (*Brachiaria*) tropical forage grasses (Tomaszewska et al., 2021a, b), apomicts at all ploidies produce seeds as there is no meiosis, nor necessity for restitution of diploid pairing behaviour. Although there is at best weak evidence that polyploid plants as a group outperform diploids either in wild or agricultural settings (particularly where selection is strong enough to remove mildly deleterious mutations), there are many prospects to exploit polyploidy systematically using additional gene alleles and increased genetic variability which enhance adaptability (Brits et al., 2003; Bhat et al., 2005; Kharrat-Souissi et al., 2014). *Cenchrus ciliaris* has a genome size of approximately 3,000 Mbp in the tetraploid species, 50% larger than the 1,971 Mbp reported for the genome of *C. purpureus* (Elephant or Napier grass; 2*n*=4*×*=28) and the more distantly related but smaller 1,580 Mbp *C. americanus* (2*n*=2*×*=14; syn. *Pennisetum americanum*) genome (Yan et al., 2021). A major proportion of plant genomes consists of repeated sequences such as ribosomal DNAs, tandem repeats, satellites, and transposable elements which play an important role in shaping of genome organization (Heslop-Harrison and Schwarzacher, 2011; Macas et al., 2011; Biscotti et al., 2015). Yan et al. (2021) reported the composition of the *C. purpureus* genome as comprising 66.32% repetitive sequences, similar to the proportion of repeats in *C. americanus* (77.2%). Ty3-*gypsy* and Ty1-*copia* are two super-families of LTR-retroelements widely distributed at the centromeric region of chromosomes in many members of Poaceae and terminal heterochromatic chromosomal regions in *Allium cepa*, respectively (Kumar et al., 1997).

Members of the major groups of repetitive elements, tandemly repeated satellites and transposable elements (Class I retrotransposons and Class II DNA transposons) differ in their abundance and genomic distribution in the genome. Tandemly repeated motifs may be located in sub-terminal, intercalary or centromeric regions of chromosomes, or, apart from the 5S and 45S rDNA arrays, not be abundant. In contrast, all plants have relatively abundant retrotransposons, often with a dispersed distribution (Heslop-Harrison et al., 1997), often dispersed throughout the euchromatin but absent from specialized regions such as heterochromatin, centromere, telomeres and nuclear organization regions (NOR). Retrotransposon families may also be restricted to characteristic genomic locations. In *Arabidopsis thaliana* many Ty3-*gypsy* retroelements are present in pericentromeric heterochromatin region (Brandes et al., 1997b). Centromeric retroelements in maize (CRM) is a lineage of Ty3-*gypsy* retroelements (Jin et al., 2004), and the diploid oat *Avena longiglumis* (2*n*=2*×*=14; 3,850 Gb genome size) has specific LTR retrotransposons located at the centromeres (Liu et al., 2022), in all cases with other retroelement families dispersed along all chromosomes. Some of these repetitive elements may have a role in centromere functionality, although they may also accumulate in regions of the genome where there are few genes.

The activity of retroelements is modulated by epigenetic mechanism through DNA methylation (Slotkin and Martienssen, 2007; Le et al., 2015), where methyl group is covalently added to the fifth carbon of cytosine residues to form 5-methylcytosine (5-mC) (Moore et al., 2013). Expression profile of retroelements can be correlated with the differential methylation pattern (Castilho et al., 1999). DNA methylation is mainly distributed in heterochromatic regions in genome that are rich in repetitive sequences consisting of retroelements (Vining et al., 2012). Comparison of methylome between the tissues or of different stages can be studied to know genes regulated through methylation and the pathways involved in response to stress and developmental conditions (Li et al., 2020). Whole genome methylation analysis can reveal the distribution pattern of methylation in the genome and the associated genes or region. In *A. thaliana*, genome-wide methylation analysis found that a third of genes are methylated within their transcribed regions while in rice about 16% of genes are enriched with methylation (Zilberman et al., 2007; He et al., 2010; Vining et al., 2012). Genome methylation analysis in *Populus trichocarpa* also indicated that Ty3-*gypsy* retroelements which are abundant in centromeric, pericentromeric and heterochromatic regions in plants are enriched with DNA methylation (Douglas and Di Fazio, 2010; Vining et al., 2012). Therefore, the methylation status of retroelements in the genome can be assessed by combining the physical mapping of retroelements through fluorescent *in situ* hybridization (FISH) with assessment of 5-methylcytosine through immunostaining which enables us to understand the genome-wide distribution pattern of retroelements and their association with the methylation.

In this study, we aimed to characterize the nature of repetitive DNA in *Cenchrus ciliaris*, investigate the genomic organization, including tandemly repeated satellite DNA and LTR retroelements, and examine genome wide methylation patterns in *C. ciliaris*, aiming to characterize features related to its contribution to the genome, evolution and possible variation in the genomes.

## Materials and Methods

### Fixation of plant material

Seeds of *Cenchrus ciliaris* (CcA7-5) were germinated in Petri dishes at 25° C for three days and root tips were collected. Seedling root tips were pretreated in alpha-bromonaphthalene for two hours to accumulate metaphases and then fixed in ethanol: glacial acetic acid (3:1) solution and stored at 4° C (Schwarzacher and Heslop-Harrison, 2000).

### Preparation of chromosome spreads

Root tips were washed twice in enzyme buffer (10 mM citric acid/sodium citrate) for 15 min and then digested with enzyme solution (20 U mL^−1^ cellulase, 10 U mL^−1^ Onozuka RS cellulase, and 20 U mL^−1^ pectinase in enzyme buffer) for 45 min at 37° C. After digestion, root-tips were again washed in enzyme buffer. The meristematic tissues were dissected and squashed in 60 % acetic acid under coverslip by using light thumb pressure. Slides were frozen on dry-ice for 5-10 min and then coverslips were removed with a razor blade. Slides were air dried and used for fluorescent *in situ* hybridization.

### Isolation of genomic DNA and designing of probes for repetitive sequences

Genomic DNA was isolated from the apomictic genotype *C. ciliaris* (CcA7-5) using the Qiagen DNeasy Plant mini kit. Primers were designed from conserved regions of the reverse transcriptase domain of LTR retroelements (Ty1-*copia* and Ty3-*gypsy*) and 45S and 5S ribosomal DNA genes using BLAST search to identify the conserved region (Sayers et al., 2019; Table 1). Whole genome sequencing data from a Chinese variety of *C. ciliaris* (deposited in SRA Sequence Read Archive under accession SRR8666664, Illumina HiSeq 2500; Nevill et al., 2020) were downloaded and used to discover the most abundant repetitive DNA sequences. Similarity-based clustering, repeat identification and classification of a subset of raw reads were performed using RepeatExplorer2 (Novak et al., 2013) after quality-trimming, excluding overlapping read pairs, and interlacing. The extracted clusters were analysed by BLAST search to check for repeat identification (Sayers et al., 2019). The primers were designed (Table 1) using Primer3 (Rozen and Skaletsky, 1999). Conserved regions were amplified from genomic leaf DNA in a standard polymerase chain reaction (PCR) using specific primers synthesized commercially (Sigma). PCR products were labeled with biotin-16-dUTP or digoxigenin-11-dUTP using BioPrime CGH array labelling kit (Invitrogen), and then purified using BioPrime Purification Module (Invitrogen).

**Table 1.**
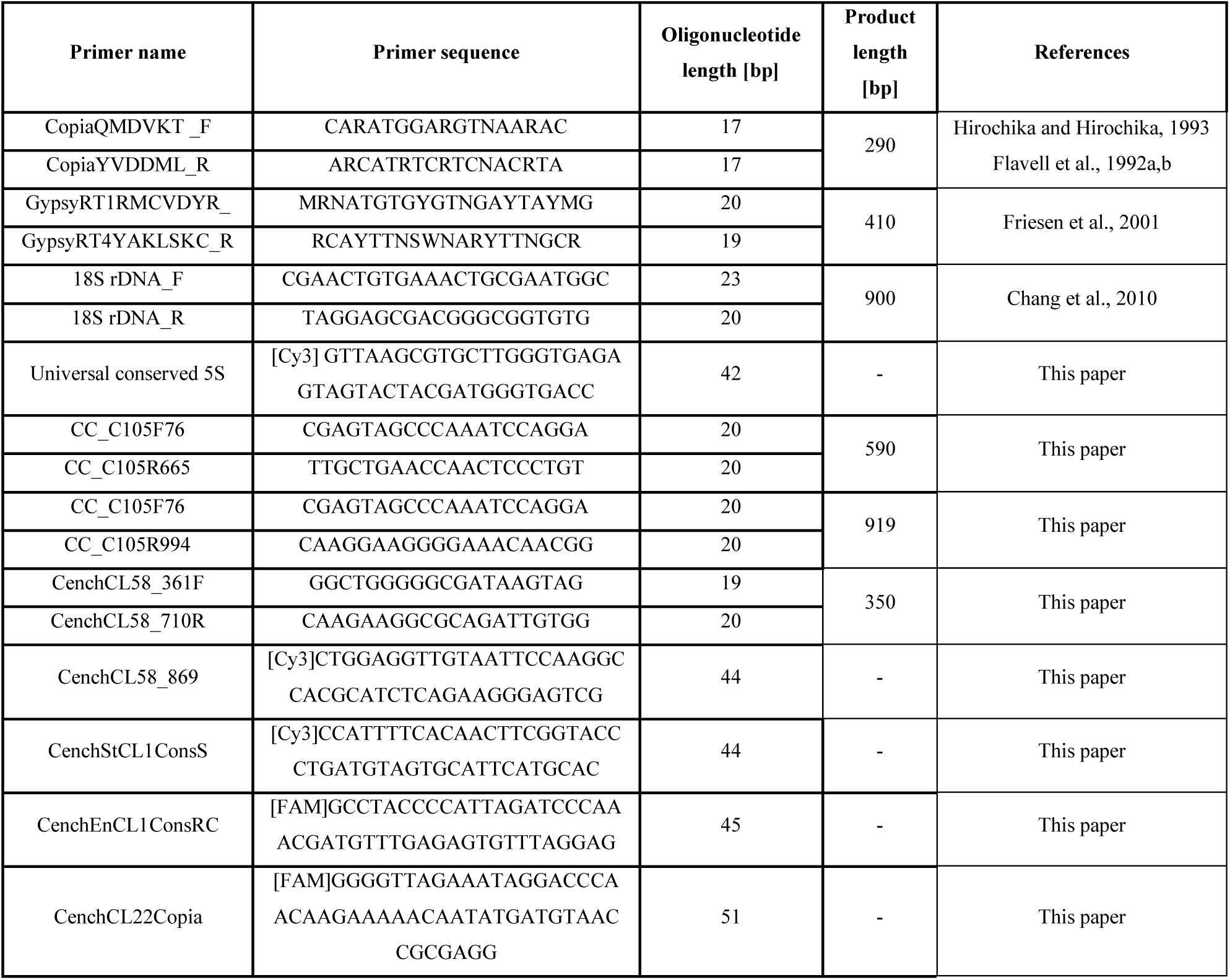
List of PCR primers and commercial oligonucleotide probes used in the study.

Oligonucleotide probes were synthesized incorporating fluorescent labels (Table 1; Sigma).

### Fluorescence *in situ* hybridization

Slides were re-fixed in ethanol:glacial acetic acid (3:1) for 30 min, dehydrated in 100% ethanol twice for 5 min and air dried. Slides were treated with RNase (100µg/ml) in 2X SSC (for 1 hr at 37° C in a humid chamber and then washed twice in 2X SSC for 5 min at room temperature.

Then slides were incubated in 0.01M HCL for 2 min, treated with pepsin (5 µg mL^-1^) in 0.01M HCl for 10 min at 37 °C in humid chamber, rinsed in distilled water for 1 min and washed in 2X SSC for 5 min. Slides were re-fixed in 4% formaldehyde at room temperature for 10 min, washed in 2X SSC for 2 min and then in 2X SSC for 10 min. Slides were then dehydrated in an ethanol series (70%, 85% and 100% ethanol, 2 min each).

Hybridization mixture consisted of probes (2 ng µL^-1^ each), 50% formamide (v/v), 2X SSC, 10% dextran sulphate (v/v), 200 ng µL^-1^ salmon sperm DNA, 0.125mM EDTA and 0.125% SDS for total volume of 40 µL was denatured at 83° C for 10 min and then cooled down on ice for 10 min. The hybridization mixture was then added to slides and placed in a hybridization oven at 73° C for 6 min to denature probe and chromosomes and then cooled to 37° C and left at that temperature overnight for hybridization.

Post hybridization washes were performed to remove unincorporated and weakly hybridized probe. Slides were washed twice in 2X SSC for 2 min at 42° C and then washed in 0.1X SSC at 42° C for 2 min and then for 10 min at the same temperature before a final wash in 2X SSC for 5 min at room temperature. To detect probes, slides were incubated in detection buffer (4X SSC containing 0.2% Tween 20) for 5 min, blocked by incubation with 5% BSA (bovine serum albumin) in detection buffer for 10 min at 37° C in humid chamber. Hybridization sites were detected by incubation with streptavidin conjugated to Alexa 594-conjugated streptavidin antibody or FITC (fluorescein isothiocyanate)-conjugated anti-digoxigenin antibody in detection buffer with 5% BSA for 1 hr at 37° C in humid chamber. Slides were washed in detection buffer thrice at 40° C for 10 min each. Chromosomes were then counterstained with DAPI (4’,6-diamidino-2-phenylindole) in antifade solution (AF1, Citifluor).

### Methylation analysis using anti-5-methylcytosine through immunostaining

Immunostaining was performed to visualize how whole-genome cytosine methylation is distributed in different chromosomal regions of *C. ciliaris* genome. Slides were pre-treated with RNase and pepsin, dehydrated in ethanol series and air dried according to FISH protocol. Slides were then blocked by 1% BSA (w/v) in 1X PBS buffer/0.5% Tween 20 for 30 min at room temperature and incubated with primary anti-5-methylcytosine antibody (anti-5-mC; 1:500 in 1X PBS buffer/0.5% Tween 20) for 1 hr at 37° C in a humid chamber. Slides were then washed in 1X PBS buffer/0.5% Tween 20 twice for 5 min each at room temperature and incubated with Alexa 488-conjugated goat anti-mouse secondary antibody (1:500) in 1X PBS buffer/0.5% Tween 20. Slides were then washed twice in 1X PBS buffer/0.5% Tween 20 for 5 min at room temperature and stained with DAPI in antifade solution (AF1, Citifluor).

### Photography and imaging

Slides were examined with a Nikon Eclipse 80i epifluorescence microscope equipped with DS-QiMc monochromatic camera, and NIS-Elements v.2.34 software (Nikon). Images were analysed using Adobe Photoshop software.

## Results

### Chromosomes of *Cenchrus ciliaris*

Most plants of the apomictic tetraploid *C. ciliaris* were 2*n*=4*×*=36 but one hyperploid 2*n*=4*×*=39 plant was found (**Fig. 1A**); all chromosomes were submetacentric and of similar length. Three major signals of 5S rDNA at intercalary regions were present (**Fig. 1B**). The 45S rDNA probe located at four sub-terminal regions (**Fig. 1C, D arrows**), also visible as the satellite at the nucleolar organizing region (NORs by DAPI staining).

**Figure 1.**
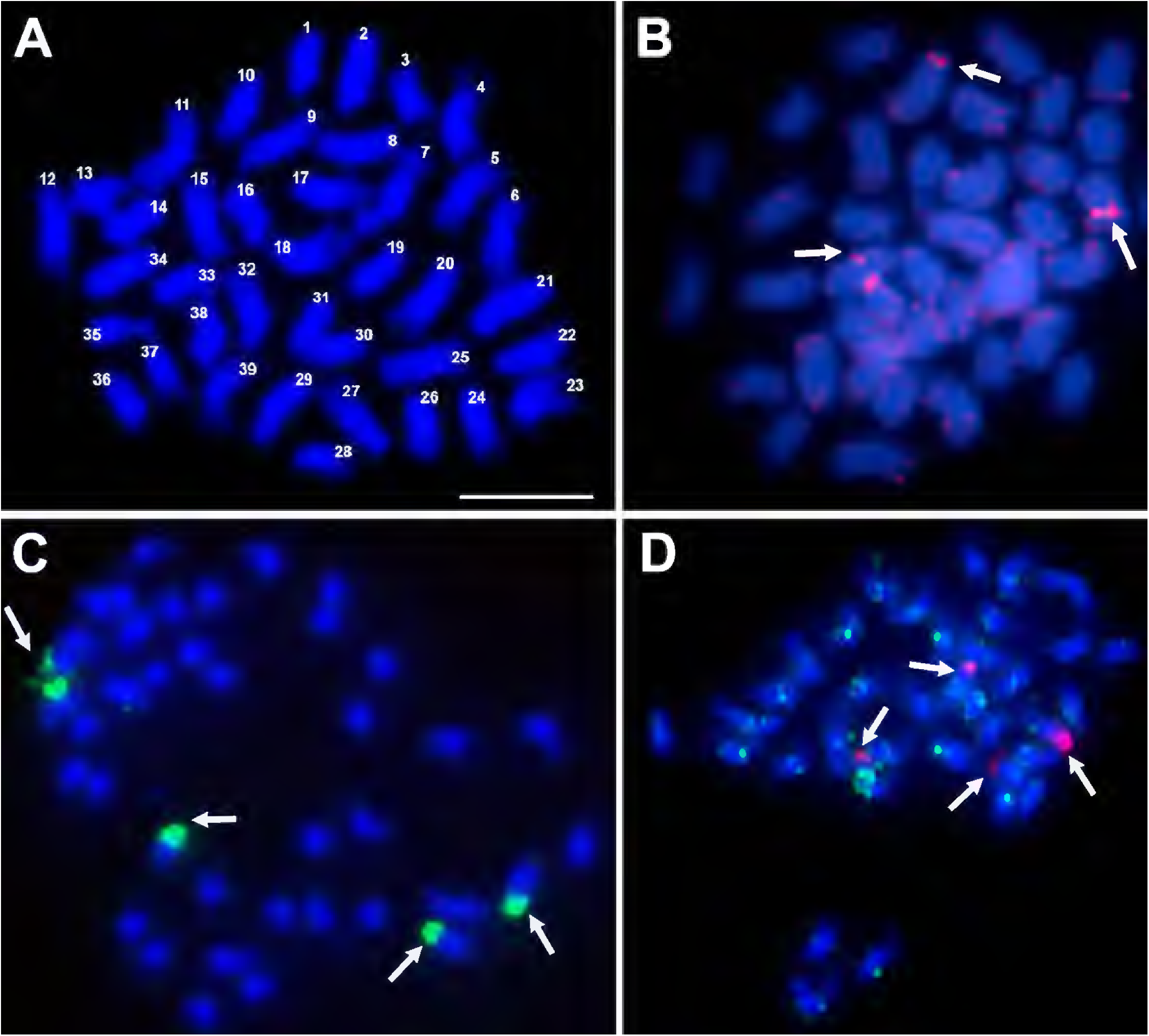
Number of chromosomes and rDNA sites in *Cenchrus ciliaris*. (A) Metaphase spread of *C. ciliaris* after DAPI staining (blue) showing a chromosome count of 2*n*=4×+3=39. (B) Metaphase chromosomes labeled with 5S rDNA fluorescing red showing three major signals (arrows). (C) Fluorescent *in situ* hybridization with 45S rDNA probe fluorescing green (arrows). (D) 45S rDNA probe (red) showing four major hybridization sites and Gy105 probe (green) hybridizing to parts of most chromosomes. Scale bar = 10μm in (A)

The degenerate PCR product used as a ‘universal’ Ty1-*copia* **(Table 1)** probe labelled broad regions of chromosomes, with sites often overlapping or spanning centromeric DAPI positive heterochromatin (**Fig 2A i**). Along with the centromeric regions, arms of most chromosomes were labeled with Ty3-*gypsy*, but most distal ends lacked signal (**Fig 2B i**). Probe Gy105 (**Table 1**), designed from a sequence including a Ty3-*gypsy* element ED545466 identified in a BAC by Conner et al. (2008) (**Fig. 1D**, green signal) labelled regions near centromeres on most but not all chromosomes.

**Figure 2.**
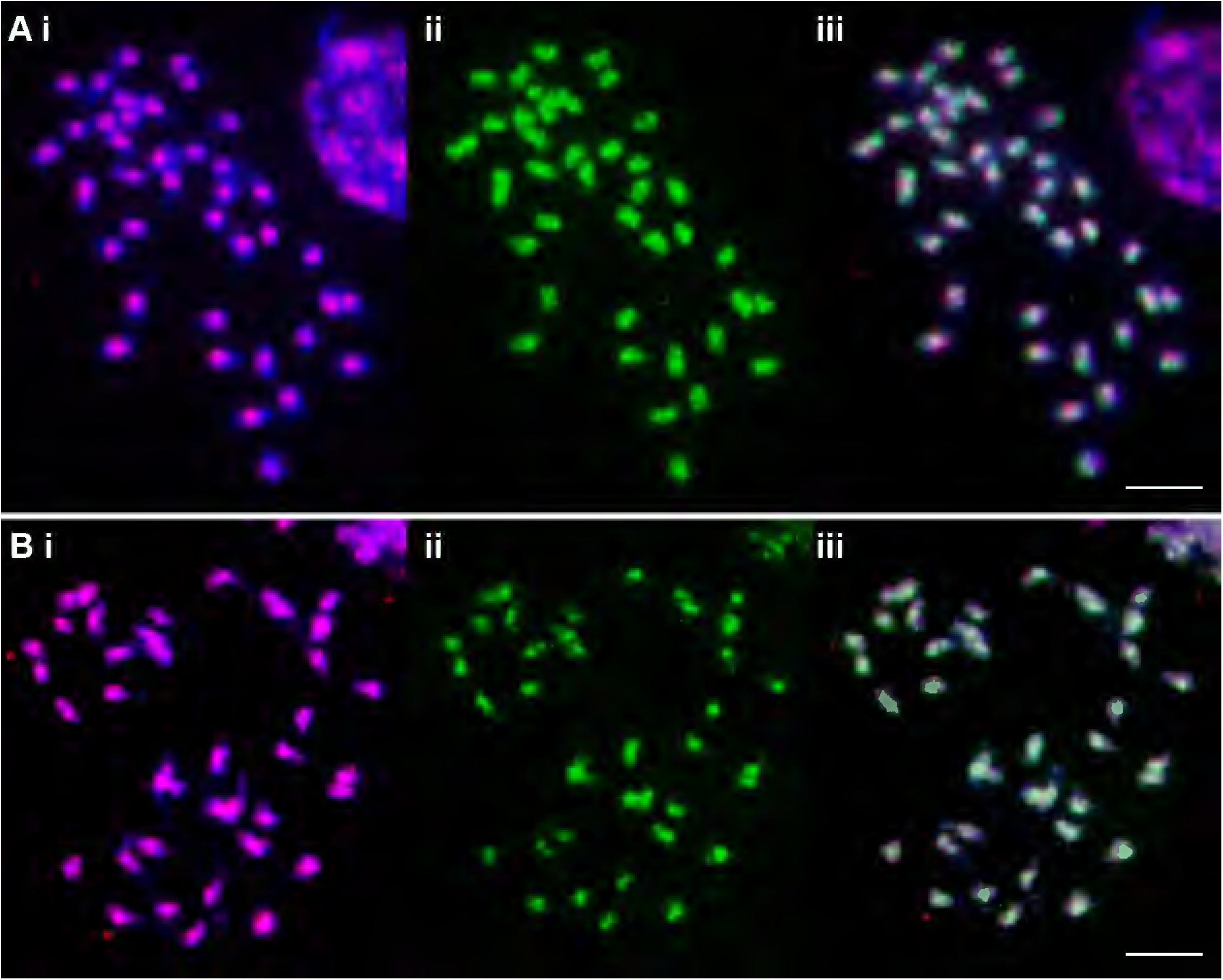
Metaphase chromosomes of *C. ciliaris* (2*n*=4×+3=39) after FISH (red fluorescing, i) and immuno-staining with anti-5-methylcytosine (green fluorescing, ii) and as overlap image (iii). (A) The ‘universal’ Ty1-*copia* probe shows concentrated signal around the centromeres of all chromosomes while 5-mC signal is seen overlapping them but also further along the chromosome arms. 5-mC strength is variable between chromosomes with most showing a small gap at the centromeres. (B) The ‘universal’ Ty3-*gypsy* probe shows a dispersed signal over the chromosome arms, but is absent from the most distal parts. 5-mC signal covers most Ty3-gypsy signal but is weak or absent in some cases (arrows). Scale bar = 10μm

Immunostaining with anti-5-methylcytosine antibody showed genome-wide methylation patterns in apomictic *C. ciliaris* along chromosomes, with variability in patterns observed. About a third of chromosome arms were more weakly labelled particularly at the ends of the arms, most centromeric regions showed small gaps, and a few chromosomes were strongly labelled (**Figs. 2A ii** and **2B ii**).

Chromosomal locations of the anti-5-methylcytosine antibody were compared with retroelement distribution by sequential labelling of the same metaphase preparations. Anti-5-mC clusters overlapped the centromeric Ty1-*copia* probe signals of most of the chromosomes (**Fig. 2A iii**). Immunostaining of chromosomal spread previously probed with Ty3-*gypsy* retrotransposons showed methylation signals of anti-5-mC overlapped with Ty3-*gypsy* retroelements on most of pericentromeric and centromeric regions of chromosomes. However, some Ty3-*gypsy* signals on the arms of chromosomes did not overlap with anti-5-mC signals (**Fig. 2B iii; arrows**). Along chromosome arms, anti-5-mC signal was generally more widespread than either retroelement.

### Chromosomal organization of highly abundant repetitive DNA motifs

Graph-based clustering (RepeatExplorer) of survey sequence of *Cenchrus ciliaris* (SRR8666664; Nevill et al., 2020) identified highly abundant repetitive DNA motifs in the genome (**Fig. 3A**).

**Figure 3.**
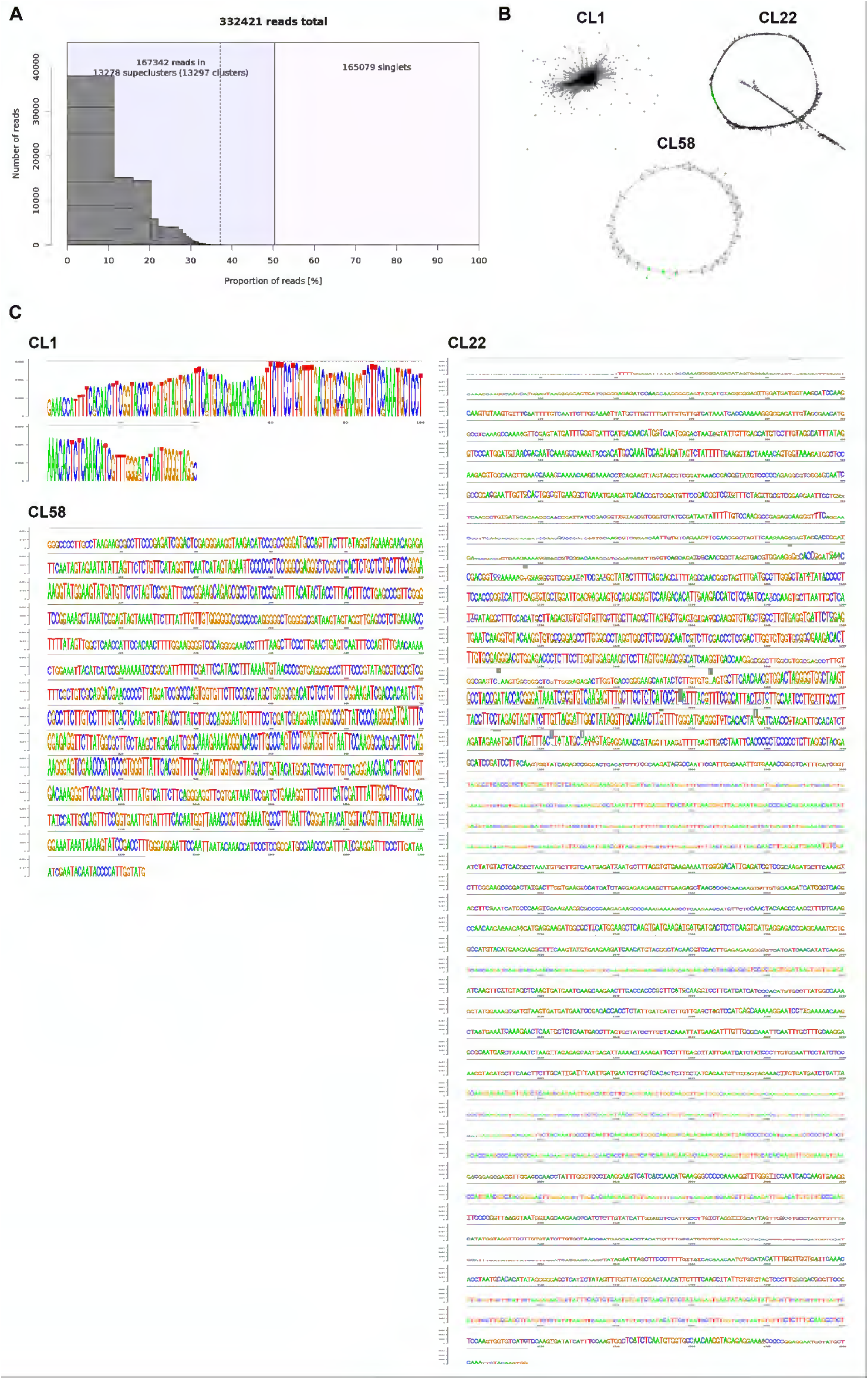
Consensus sequences used to design primers and probes for fluorescence *in situ* hybridization. A) RepeatExplorer graph showing number of raw sequencing reads forming abundant clusters; about 50% of the sequences form clusters, with less than 20 motifs representing a third of the genome. B) Graphical 2D projection of the structure of three graph-based clusters found in *Cenchrus ciliaris* genome using RepeatExplorer. Each node represents one of the sequence reads. The placement of the nodes reflects sequence similarity and overlaps. Cluster1 and Cluster58 are tandem repeats while cluster22 is a LTR-copia element. C) Sequence LOGO representations showing lengths of repetitive motifs and conservation of sequences. Cluster1 is 140bp tandem repeat; Cluster58 consensus is 1326bp and Cluster22 4807bp.

Cluster1 (**Supplementary Data Table S1; Fig. 3B**), accounted for 2.5% of the *C. ciliaris* genome, and is a putative low-confidence satellite of 140bp monomer length (**Fig. 3C**), part of which is homologous to a similarly sized satellite reported in *Setaria viridis* (Thielen et al., 2020). Two oligonucleotide probes (44 and 45 bp long) were designed from two different parts of this sequence: CenchStCL1ConsS and CenchEnCL1ConsRC (**Table 1**). Cluster58 (**Supplementary Data Table S1; Fig. 3B, C**) is identified as a satellite with a repeat unit of 1326bp that represents a much smaller proportion, 0.061% of the genome, and PCR primers and a 44bp probe were designed for this sequence (**Table 1**). Cluster22 (**Supplementary Data Table S1; Fig. 3B, C**) is identified as a 4807bp Ty1-*copia* retrotransposon, and the oligonucleotide probe CenchCL22Copia was designed to the RNaseH domain (RC) from this sequence (**Table 1**). The probes were localized on metaphase chromosomes of *C. ciliaris* using fluorescence *in situ* hybridization.

CenchEnCL1ConsRC probe gave strong signals mostly in pericentromeric regions of about twenty chromosomes from each metaphase, and the remaining chromosomes showed very weak signals (**Fig. 4A**). Twenty very strong signals of CenchStCL1ConsS probe were also detected in pericentromeric regions of chromosomes, and some minor signals, mostly intercalary, were visible (**Fig. 4B**). CenchStCL1ConsS collocalized with CenchCL22Copia signals (**Fig. 4C**). Both probes are good chromosome markers distinguishing about half the chromosomes. Amplified CenchCL58 probe and commercially synthesized CenchCL58_869 showed dispersed signals along all chromosomes from metaphase (**Fig. 4D**). Stronger signals of CenchCL58_869 probe collocalized with CenchEnCL1ConsRC (**Fig. 4E**), with 19 weaker and 20 stronger centromeric sites.

**Figure 4.**
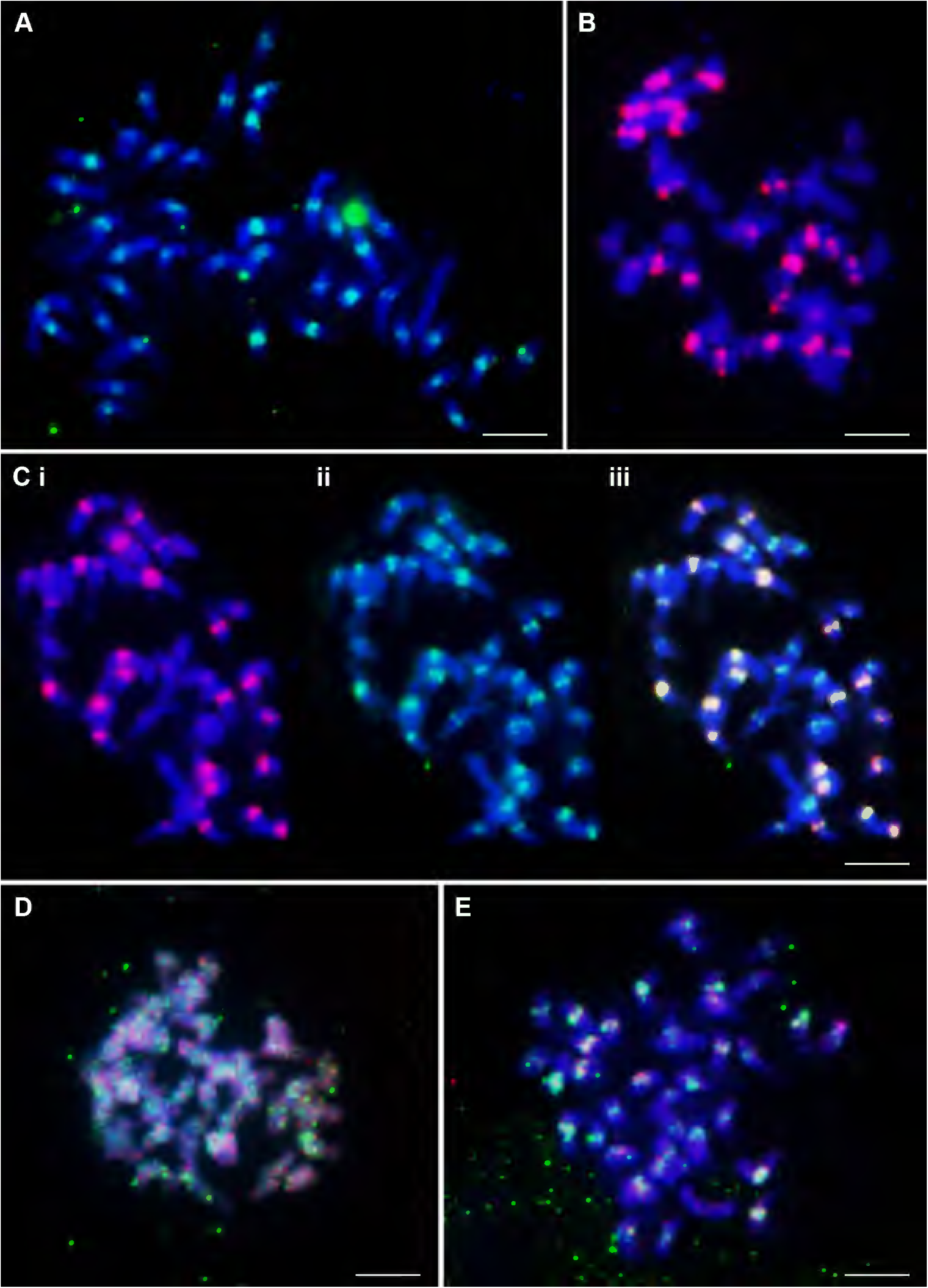
Distribution of highly abundant repetitive DNA motifs on chromosomes of *C. ciliaris* (2*n*=4×+3=39). (A) Tandem repeat CenchEnCL1ConsRC probe labeled commercially with FAM (green) showed strong signals in pericentromeric regions of about half of the chromosomes. The remaining chromosomes showed weak signals. (B) Twenty very strong and 4 weaker red signals of Tandem Repeat CenchStCL1ConsS probe labeled commercially with Cy3 (red). (C) Three pictures of the same metaphase showing collocalization of Cluster1 and Cluster22 signals: (i) CenchStCL1ConsS (labeled with Cy3; red) showing signal on about half of the chromosomes, (ii) CenchCL22Copia (labeled with FAM; green) with signal on all chromosomes, (iii) merged. (D) Metaphase chromosomes showing dispersed signals of amplified CenchCL58 probe (detected with FITC; green) and commercially synthesized CenchCL58_869 (Cy3, red). (E) Dispersed signals of CenchCL58_869 probe (Cy3, red) along chromosomes. Some stronger signals collocalize with CenchEnCL1ConsRC (FAM; green). Scale bar = 10μm

## Discussion

### Aneuploidy and polyploidy in *Cenchrus ciliaris* – consequences for forage grasses

*C. ciliaris* is an allotetraploid with chromosome number 2*n*=4*×*=36 (Akiyama et al., 2011) and a primary base number of *×*=9. In this study, we reported a new chromosomal count of *C. ciliaris* with 2*n*=4*×*+3=39. Aneuploidy in *C. ciliaris* has been widely reported (Visser et al., 1998), with Burson et al. (2012) finding 99 aneuploids from 568 accessions, with different chromosome numbers from 2*n*=18 to 2*n*=56 (Vij and Chaudhary, 1981). Hyperploidy or monosomic addition lines are well-known in tetraploids and hexaploids in many Poaceae members, including *C. ciliaris* (2*n*=6*×*+1=55). Meiotic irregularities have been widely reported in *C. ciliaris*, presumably because of the close genomic relationships in the polyploid (Visser et al., 1998; Burson et al., 2012). Where seed production is continued through apomixis, the predominant pathway in *C. ciliaris*, there is low selective pressure for euploids. Polyploidization has significant conservational and environmental consequence where one species can become invasive due to adaptation of a species in a new locality and can also change reproductive biology (Thompson and Pellmyr, 1992; John et al., 2004). Thus, characterization of the genome constitution of apomictic *C. ciliaris* is important both genetically and environmentally.

Aneuploid and polyploid cytotypes are widely distributed within tropical forage grasses, including species of the genera *Cenchrus, Urochloa, Panicum, Pennisetum*, and *Paspalum* (Carloni-Jarrys et al., 2018; Worthington et al., 2019; Tomaszewska et al. 2021a, b). Polyploids can arise from spontaneous chromosome doubling following non-disjunction after meiotic or mitotic divisions (Gallo et al., 2007; Burson et al., 2012; Worthington et al., 2019), and often involve fertilization with unreduced (2*n*) gametes. Many species, such as those in the Triticeae tribe, have mechanisms that control chromosome pairing (Luo et al., 1996; Sepsi et al. 2017) and restore diploid meiotic behaviour and fertilization, with elimination of unpaired chromosomes. In contrast, where seed is produced through apomixis, many ploidies and aneuploid lines can be maintained and propagated unchanged (maintaining also any heterozygosity or heterosis in the maternal genotype; Lippman and Zamir, 2007). Apomixis enables propagation of outstanding heterozygous genotypes without vegetative propagation or seeds, which would be especially valuable for domesticated grass breeding (Ozias-Akins and van Dijk, 2007). Along with the 2*n*=4*×*+3=39 *C. ciliaris* reported here, there are many examples of aneuploids in apomicts at various ploidies such as *Urochloa humidicola*, where an aneuploid accession with 2*n*=8*×*+2 (or 9*×*−4) = 50 has been found (Tomaszewska et al., 2021a). However, unlike tropical forage grasses, including segmental allopolyploids (Jessup et al., 2003; Worthington et al., 2019; Tomaszewska et al. 2021b), cereal grain-crop species tend to exhibit variation in levels of ploidy and relatively low genome homology, which constitute a barrier for crossbreeding with apomicts and make it impossible to achieve agronomically acceptable genotypes. Identification of loci that firstly control heterotic phenotypes and secondly parthenogenesis or apomixis in tropical forage grasses is important for forage grass improvement, and further work may lead to suitable systems to exploit apomixis in grain crops, regarded as a significant challenge for wheat and rice (Dresselhaus et al., 2001; Conner and Ozias-Akins, 2017; Rathore et al. 2020), thus fixing heterozygosity and removing the requirement for sexual seed production.

### Chromosomal localization of highly abundant repetitive DNA motifs

Various approaches have been used to measure the abundance and genome organization of repetitive DNA in the plant genome, and assemblies from short reads will never give the amount present because of collapse of similar repeats during assembly (e.g. Yan et al., 2021). Graph-based clustering of unassembled raw reads, using RepeatExplorer (Novak et al., 2013) is proving to be a reference-free approach to characterize the number and abundance of all repeat families in a genome. Analysis of the frequency of k-mers – oligonucleotide sequences k bases long, where k is typically 16 to 64bp – has also proved a robust method to identify repetitive motifs (Liu et al., 2019; Tomaszewska et al., 2021b).

Here, we used both specific probes developed using graph-based sequence clustering compared to ‘universal’ conserved regions of Ty1-*copia* and Ty3-*gypsy* retroelements. Similarly to *Urochloa* tropical forage grasses, where genomic DNA and retroelements used as probes showed no substantial differences between genomes (Santos et al., 2015; Tomaszewska et al., 2021b), parallel universal retroelement repeats used in this study were relatively equally abundant over the whole *Cenchrus ciliaris* genome (see **Fig. 3**). The Gy105 probe (see **Fig. 1D**) labelled some chromosomes more weakly. In contrast, Cluster1 (putative satellite) and Cluster22 (LTR-copia) probes showed clear differential labelling of about half the chromosomes (see **Fig. 4**), and we suggest differentiate the two ancestral genomes in the allopolyploid *C. ciliaris* having different retroelement (and/or satellite) sequence variants amplified before the genome came together in the tetraploid. As might be expected, the ‘universal’ retroelement probes amplified abundant sequences from both genomes and did not show any genome differentiation (see **Fig. 2**).

### Methylation pattern and major repeat distribution

The genome distribution and organization of retroelements is important for understanding retroelements dynamics, movement or amplification, chromosomal structure, and the evolutionary processes. In this study, centromeric DAPI-staining bands (corresponding to constitutive heterochromatin; Siljak-Yakovlev et al., 2002) collocated with Ty1-*copia* probes, suggesting accumulation of these retroelements at heterochromatic regions. Both Ty1-*copia* and Ty3-*gypsy* tended to cluster at specific regions on most of chromosomes (along with some intercalary and dispersed sites) contrasting with other species (Brandes et al., 1997a) while Ty1-*copia* was more uniformly distributed on the euchromatin regions in the genome. Whether the non-uniform distributions of retroelements is due to insertion-site preference in the genome (Belyayev et al., 2001), perhaps of particular element subfamilies, or purging from genome regions, will need further study.

A combination of *in situ* hybridization with immunostaining using anti-5-mC allows correlation of the distribution of DNA methylation and sequence location across the genome (Sepsi et al., 2014). 5-mC has been extensively studied for its role in regulation of gene expression, genome imprinting, and suppression of transposable elements (Zhang et al., 2010). In this study, anti-5-methylcytosine immunofluorescence overlapped with most of the Ty1-*copia* elements and Ty3-*gypsy*, showing their location predominantly in methylated genomic regions associated with lower transcriptional activity. With the majority of 5-mC signal associated with retrotransposon-rich regions, there was no further differentiation of groups of chromosomes, and, in particular, no indication of differences between ancestral genomes in the polyploid. In contrast, differential methylation of parental genomes has been observed in intraspecific *A. thaliana* hybrids and implicated in heterosis as a key epigenetic factor (Lauss et al. 2018). However, in rice tetraploid hybrids, Li et al. (2019) reported overall methylome stability with regional variation of cytosine methylation states, in part associated with gene expression changes. Methylation repatterning, often stimulated by hybridization and/or polyploidization, can potentially lead to increased genetic variation through facilitating somatic recombination, which could be adaptive in apomictic lineages (Verhoeven et al., 2010). Changes in methylation pattern were observed both in the newly formed and in the established apomicts of *Taraxacum*, suggesting that considerable methylation variation can increase over short evolutionary time scales.

## Author Contributions

Conceptualization, all authors.

Funding acquisition, PR, VB, PHH and PT.

Investigation and Methodology, PR, TS and PT

Supervision, VB and PHH

Writing original draft review & editing, all authors.

All authors have read and agreed to the published version of the manuscript.

## Funding acknowledgement

P.R. was supported under the Newton Bhabha Ph. D placement programme fellowship coordinated by Department of Biotechnology, India and British Council, United Kingdom (BTIIN/UKJDBT-BC/2017-18). This work was supported under the RCUK-CIAT Newton-Caldas Initiative “Exploiting biodiversity in *Brachiaria* and *Panicum* tropical forage grasses using genetics to improve livelihoods and sustainability”, with funding from UK’s Official Development Assistance Newton Fund awarded by UK Biotechnology and Biological Sciences Research Council (BB/R022828/1). P.T. has received support from the European Union’s Horizon 2020 research and innovation programme under the Marie Sklodowska-Curie grant agreements No 844564 and No 101006417 for analysis of polyploid chromosomal evolution.

## Data availability statement

Previously published sequence data is available under accession SRR8666664 (Nevill et al. 2020). Results of the RepeatExplorer2 analysis of these sequence reads are archived on Figshare at http://dx.doi.org/10.25392/leicester.data.19798966.

---

**Supplementary Data Table S1.**
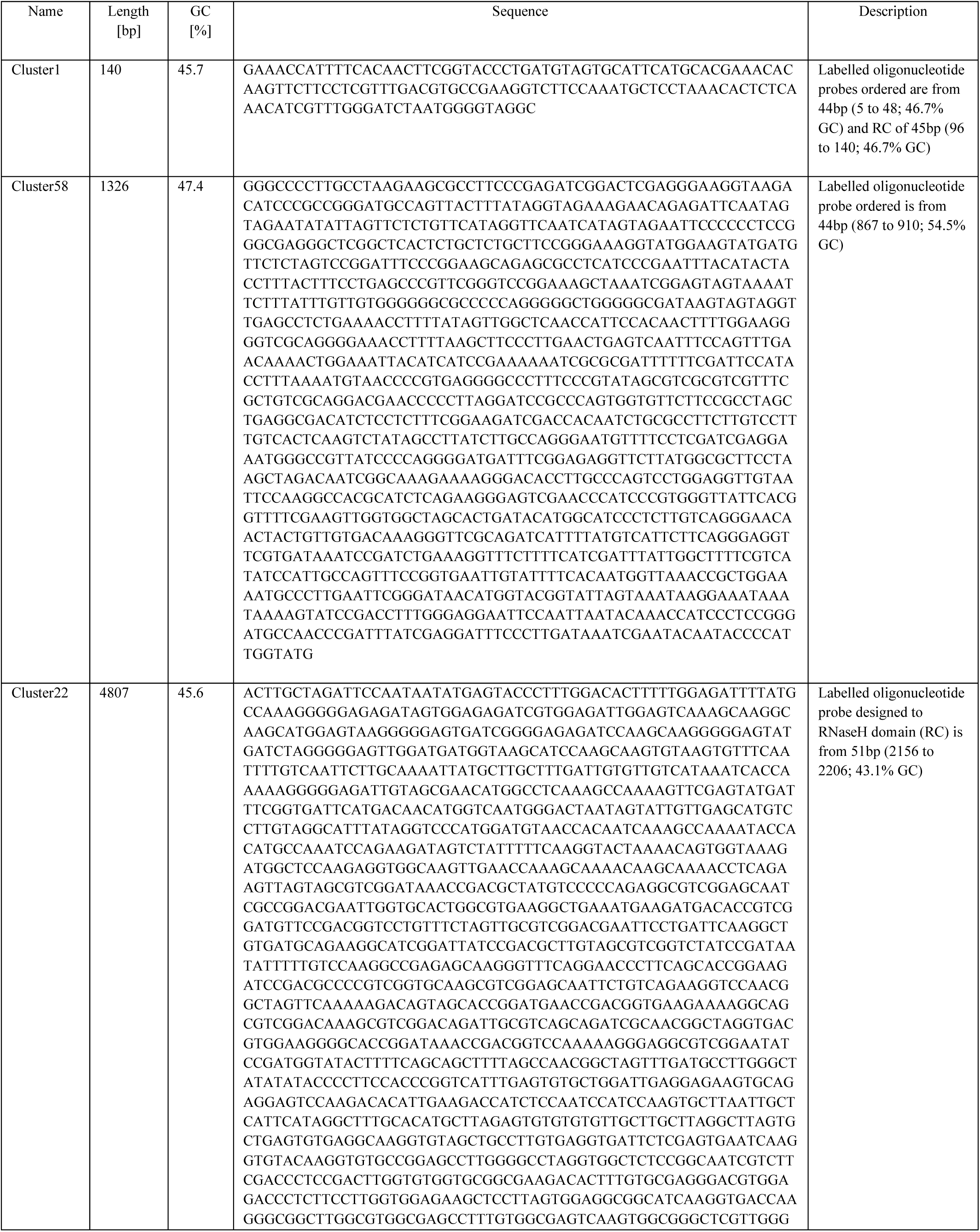
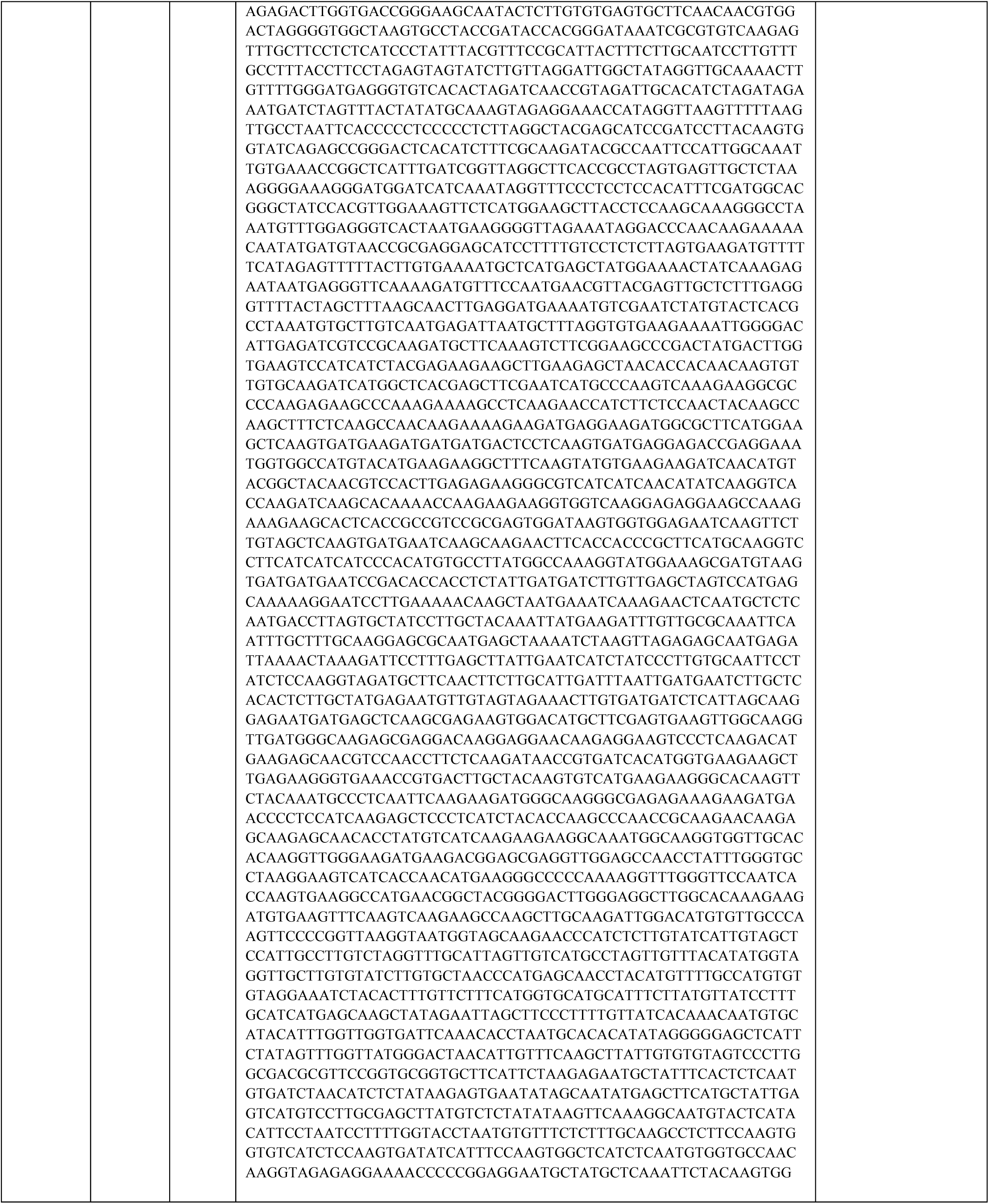
List of the most abundant clusters extracted from the Chinese variety of *Cenchrus ciliaris* SRR8666664 using RepeatExplorer.

